# ECHO: a nanopore sequencing-based workflow for (epi)genetic profiling of the human repeatome

**DOI:** 10.64898/2026.03.18.712618

**Authors:** B. Poggiali, L. Putzeys, J. D. Andersen, A. Vidaki

**Affiliations:** Department of Clinical Genetics, Maastricht University Medical Centre, 6229 HX Maastricht, The Netherlands; Section of Forensic Genetics, Department of Forensic Medicine, Faculty of Health and Medical Sciences, University of Copenhagen, 2100 Copenhagen, Denmark; Department of Genetics and Cell Biology, GROW Research Institute, Maastricht University, 6229 HX Maastricht, The Netherlands

## Abstract

**Summary:** The human genome is dominated by repetitive DNA, whose genetic and epigenetic variation plays a key role in gene regulation, genome stability, and disease. Recent advances in long-read sequencing now enable large-scale, haplotype-resolved, and DNA methylation-informative analysis of the human genome, including on previously inaccessible complex and repetitive regions. However, the comprehensive, simultaneous characterisation of the “human repeatome” remains challenging, largely due to the lack of comprehensive tools integrated in a single pipeline that can capture the full spectrum of variation across diverse types of DNA repeats. Here, we present ECHO, a user-friendly, Snakemake-based pipeline for the “(Epi)genomic Characterisation of Human Repetitive Elements using Oxford Nanopore Sequencing”. ECHO provides a reproducible and scalable framework for end-to-end analysis of whole-genome nanopore sequencing data, enabling integrative but also tailored (epi)genetic analyses of the human repeatome.

**Availability and implementation:** ECHO is freely available at Github: https://github.com/leenput/ECHO-pipeline, with the archived version at Zenodo: https://zenodo.org/records/19068468

## 1 Introduction

Repetitive DNA elements account for more than half of the human genome (Nurk *et al*., 2022). These include tandem repeats (TRs) that are characterized by arrays of repeated DNA sequence units, and interspersed repeats or transposable elements (TEs) that are scattered throughout the genome (Liao *et al*., 2023). Despite their abundance, repeats have remained underexplored in human genomics, primarily due to technical limitations of short-read sequencing and array-based technologies, which are poorly suited to resolve long, complex, and repetitive regions (Treangen and Salzberg, 2012). This technical barrier, among other factors, has contributed to the longstanding misconception that repetitive DNA lacks functional relevance (Ohno, 1972).

In recent years, however, this perspective has shifted, since growing evidence continues to highlight key roles of repetitive DNA in genome organization, gene regulation, evolution, and disease (Wright and Todd, 2023; Lawson *et al*., 2023; Fueyo *et al*., 2022). Specifically, TR expansions can cause neurological and developmental disorders (Depienne and Mandel, 2021), while TE-mediated genomic rearrangements and insertional mutagenesis can contribute to tumorigenesis and other complex diseases (Chuong *et al*., 2017). Beyond sequence-driven effects, repeats are tightly regulated by epigenetic mechanisms, particularly DNA methylation, to modulate transcription and preserve genome integrity. Importantly, both TRs and TEs display dynamic, context-specific DNA methylation patterns, which may vary across haplotypes, cell types, developmental stages, environmental cues, and disease states (Pappalardo and Barra, 2021; Barbé and Finkbeiner, 2022). As such, to fully understand the regulatory and functional scope of repeats at a genome-scale, we require integrated analysis of their (epi)genomic features.

More recently, the renewed momentum in repetitive DNA research has been catalysed by the advent of long-read sequencing (LRS) technologies, such as those developed by Oxford Nanopore Technologies (ONT) and Pacific Biosciences (PacBio) (Logsdon *et al*., 2020). These platforms uniquely enable genome-wide access to complex and repetitive regions, along with direct measurement of DNA methylation, at single-molecule resolution and high-accuracy (Fu *et al*., 2025). However, current LRS-based repeat genotyping tools are typically confined to specific repeat types (TRs or TEs, or their subclasses) (Ziaei Jam *et al*., 2024; De Coster *et al*., 2024; Tanudisastro *et al*., 2024; Bilgrav Saether and Eisfeldt, 2024; Chu *et al*., 2021). While most of these tools primarily focus on sequence variation, notable exceptions have demonstrated the potential of LRS to jointly profile the sequence and methylation of repeats. For example, TRGT enables TR (epi)genotyping from PacBio HiFi reads (Dolzhenko *et al*., 2024), while TLDR identifies non-reference TE insertions and supports downstream methylation analysis through a set of accompanying scripts (Ewing *et al*., 2020).

To date, repeat (epi)genotyping remains fragmented across repeat types, specialized tools, and sequencing platforms. Consequently, a unified, genome-wide framework for concurrent analysis of multiple repeat classes in terms of sequence and methylation states is still lacking. To address this for ONT data, we present “ECHO”, a comprehensive Snakemake-based pipeline for the “(Epi)genomic Characterisation of Human Repetitive Elements using Oxford Nanopore Sequencing”.

## 2 Pipeline description

### 2.1 Overview and implementation

The ECHO pipeline integrates benchmarked bioinformatics tools and custom Bash/Python scripts in an end-to-end workflow comprising two core stages: (I) ONT data preprocessing and phasing, and (II) repeatome profiling across TR and TE classes, integrating both sequence and DNA methylation variation (Figure 1A). The first phase handles raw ONT files, performs quality control (QC) and optional read filtering, followed by read alignment, variant calling, and haplotype phasing. The second phase leverages phased alignment and variant files to perform repeat (epi)genotyping of predefined loci of interest at a genome-wide or targeted manner. It quantifies haplotype-specific DNA methylation with single-CpG and region-level resolution; the latter aggregated across all CpGs within a repeat and its flanking regions. Overall, the pipeline is implemented in a Snakemake workflow management system (Köster *et al*., 2021). All software dependencies are managed via Singularity to ensure reproducibility and portability (Kurtzer *et al*., 2017). ECHO is designed to run both on unix-based high-performance computing (HPC) systems and on local servers.

**Figure 1:**
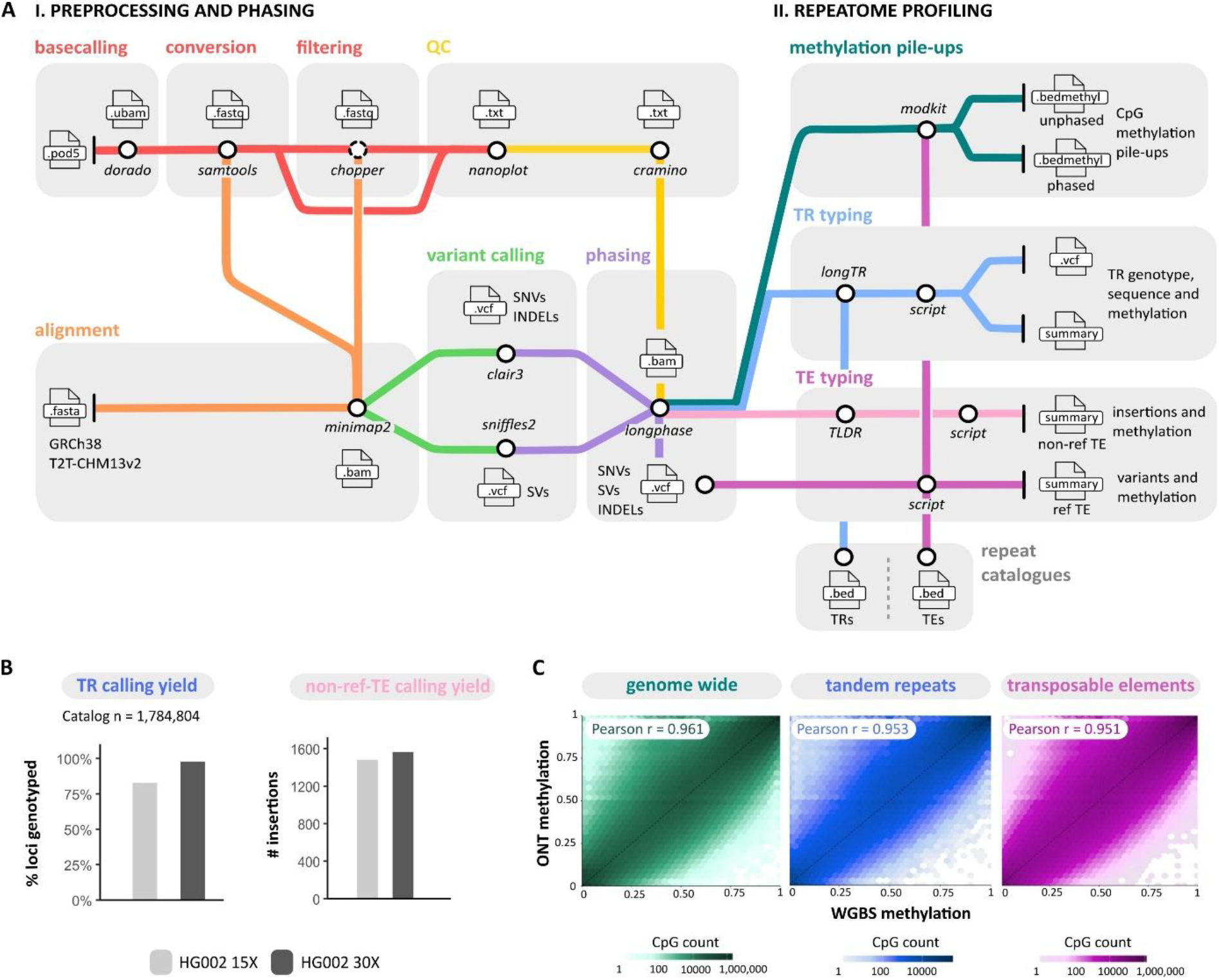
Overview and testing of the ECHO pipeline for human repeatome profiling using ONT data. **(A)** Two main modules of the pipeline: (I) preprocessing and phasing, and (II) repeatome profiling. Module I includes basecalling, QC, read filtering, alignment to a reference genome (GRCh38 or T2T-CHM13v2), variant calling, and phasing to generate haplotype-resolved BAM and VCF files. Based on user-defined repeat catalogues, module II performs TR and TE genotyping (both reference and non-reference TE insertions), along with the extraction of haplotype-specific DNA methylation at single-CpG resolution across the repeat region, as well as averaged across the repeat element and its up- and downstream flanking regions. External bioinformatic tools and custom Bash scripts are shown in italics, and key output file types are indicated at each step. (**B**) Yield of called repeat loci in HG002 across sequencing coverage (30× vs 15×). Left: percentage of TR loci successfully genotyped by LongTR relative to the genome-wide TR catalog (n = 1,784,804). Right: number of detected non-reference TE insertions by TLDR (filter = PASS). (**C**) Concordance between ECHO- and WGBS-derived DNA methylation values for the HG002 30× dataset. Hexbin (2D density) plots compare CpG methylation levels measured by ONT and WGBS genome-wide, within TRs (defined using genome-wide Adotto catalogue (English *et al*., 2025)) and TEs (defined using GRCh38 RepeatMasker annotation). Bin colour indicates the number of CpGs per hexagon. Pearson correlation coefficients (*r*) are indicated; dashed lines represent y = x. *INDEL – insertion–deletion; non-ref TE – TE not present in the reference genome; ONT – Oxford Nanopore Technologies; QC – quality control; r – Pearson correlation coefficient; ref TE – TE annotated in the reference genome; SNV – single nucleotide variant; SV – structural variant; TE – transposable element; TR – tandem repeat; WGBS – whole-genome bisulfite sequencing*.

### 2.2 Input data

As input ECHO accepts ONT sequencing data in multiple formats, including raw POD5 files, or basecalled unaligned BAM (UBAM), FASTQ, and BAM files, provided that methylation-aware ONT models have been used for basecalling. In addition, ECHO requires repeat catalogues of TR and TE target regions, defined relative to the human reference genome in use (GRCh38/T2T-CHM13v2). To facilitate standardized analyses, ECHO provides several publicly available repeat catalogues and derived subsets thereof that are reformatted for downstream compatibility, including both genome-wide and targeted panels (See Github documentation). Users may also supply custom catalogues in BED format, if preferred.

### 2.3 Preprocessing and phasing

When raw sequencing files (POD5) are provided and GPU resources are available, basecalling is performed using Dorado (v1.1.1) in methylation-aware super accuracy (SUP) mode. For UBAM input, reads are converted to FASTQ files using samtools (v1.22) with transfer of methylation tags (Li *et al*., 2009). Optional read filtering is carried out using Chopper (v0.11.0), based on user-defined thresholds for minimum read length and quality (De Coster and Rademakers, 2023). The reads are then aligned to the human reference genome (GRCh38 (default) or T2T-CHM13v2) using minimap2 (v2.30) with default settings for ONT data (-ax map-ont). QC metrics for raw, filtered, and aligned reads are obtained using Cramino (v1.1.0) and NanoPlot (v1.46.1) (De Coster and Rademakers, 2023).

After alignment, reads are processed for genome-wide variant calling. Single nucleotide variants (SNVs) and insertions-deletions (INDELs) are identified using Clair3 (v1.2.0) (Zheng *et al*., 2022), while structural variants (SVs) are detected with Sniffles2 (v2.6.3) (Smolka *et al*., 2024). LongPhase (v1.7.3) is subsequently used for co-phasing of small variants (SNVs and INDELS) and SVs in conjunction with methylation information, after which reads are haplotagged to produce a phased alignment file (Lin *et al*., 2022). Phasing QC is conducted using cramino (--phased, --karyotype) (v1.1.0), which provides relative chrX and chrY coverage metrics that are used for sex inference. The resulting phased BAM and VCF files are used for downstream repeatome analysis.

### 2.4 Repeatome profiling

#### 2.4.1 Genome-wide CpG methylation

As a first step, genome-wide CpG methylation is profiled using modkit (--cpg --ignore h) (https://github.com/nanoporetech/modkit) (v0.5.0), generating both unphased and phased methylation pileups from the phased BAM file. The resulting bedMethyl files can be used to extract methylation levels across any region in the reference genome, including for the repeats of interest (see 2.4.2 and 2.4.3).

#### 2.4.2 Tandem repeats

TR genotyping is carried out with LongTR (v1.2) using recommended ONT-optimized alignment parameters, which has demonstrated high genotyping accuracy and robustness in a recent benchmarking study (Ziaei Jam *et al*., 2024; Aliyev *et al*., 2026). LongTR takes as input the phased BAM file along with a user-defined TR catalogue specifying the target regions and generates a phased VCF output, which is subsequently processed to integrate TR genotypes with methylation information. For each TR locus, overlapping reads are extracted in a haplotype-aware manner and locally realigned to their respective TR allele sequences and flanking regions. Based on these local alignments, single-CpG and average methylation levels per TR and their flanking regions are quantified for each haplotype using modkit (--cpg --ignore h --mod-threshold m:0.8). In addition, the uTR tool (v1.0.0) is used to decompose the TR motif composition of each TR allele (Masutani *et al*., 2023). The script outputs an annotated VCF file and summary table reporting the TR allele lengths, sequences, motif structures, and methylation at three resolution levels: single-CpG within the TR, and regional averages across the TR and its flanking regions.

By default, ECHO uses a genome-wide TR catalogue adapted for compatibility with LongTR (1,784,804 loci, GRCh38), including short tandem repeats (STRs, 1-6 bp repeat units) and variable number tandem repeats (VNTRs, longer repeat units) (English, 2024). In the default pipeline configuration, downstream TR methylation analysis focuses on loci whose canonical repeat motif corresponds to an STR and contains at least one CpG site within the repeat motif (8,424 loci) to avoid computational burden. Nevertheless, ECHO supports several alternative, pre-configured TR catalogues tailored to specific applications or reference assemblies, as well as custom user-defined catalogues (see GitHub documentation).

#### 2.4.3 Transposable elements

TE analysis is separated into two components: 1) characterisation of TEs annotated in the reference genome (ref-TEs) and 2) identification and characterisation of TE insertions not present in the reference genome (non-ref-TEs). For the ref-TEs, the workflow provides catalogues derived from UCSC Genome Browser RepeatMasker tracks (Casper *et al*., 2026). By default, a complete GRCh38 TE catalogue is used, with optional support for a T2T-CHM13v2 catalogue and TE type-specific subsets (e.g. LINEs, SINEs, SVAs, helitron, DNA transposons). For each locus in the catalogue, SNVs and SVs from phased VCF files passing all variant filters (FILTER=PASS) are intersected with the ref-TE regions using BCFtools (Danecek *et al*., 2021) and bedtools (Quinlan and Hall, 2010) to quantify sequence variation per element. In parallel, single-CpG and regional methylation levels are calculated separately for each TE across its body and upstream and downstream flanking regions using phased and unphased modkit pileups, yielding allele-specific and aggregated methylation estimates, respectively. In the final output directory, a summary file reports each ref-TE with its genomic location, classification, regional methylation (phased/unphased), and counts of overlapping SNV and SVs. CpG resolutions methylation values and overlapping variants for each ref-TE region are stored in modkit pileup BED files and variant VCF files within the same directory, enabling fine-grained analysis of TE loci of interest.

In parallel, non-ref-TE insertions are identified and classified from the ONT reads using the widely adopted long-read TE detection tool TLDR (Ewing *et al*., 2020). TLDR outputs a single summary file listing all detected insertions and, for each individual TE insertion, a locally assembled TE consensus sequence (FASTA format) and a corresponding alignment file (BAM format). Based on these outputs, ECHO implements a custom bash script to filter for high-quality insertions (filter = PASS) and uses modkit to subsequently calculate single-CpG and regional methylation levels per insertion and its flanking regions. The resulting ECHO output extends the TLDR summary with regional methylation metrics for each non-ref-TE (phased/unphased), complementing the reference-based TE analysis. Detailed modkit pileup files are also retained in the output directory, enabling methylation analysis at single-CpG resolution.

## 3 Installation and usage

ECHO is distributed and maintained via a public GitHub repository, which provides detailed installation instructions, documentation, and test data. The workflow requires Snakemake and Singularity to be available on the target system. Snakemake pipeline configuration is defined through a YAML file specifying sample lists, reference genome, TR and TE catalogues, input data formats, and additional analysis parameters. Configuration files can be generated using a dedicated Python script, which also performs consistency checks and validation prior to execution. Bundled ECHO repeat catalogues are hosted on Zenodo and can be downloaded upon installation to ensure consistency with the pipeline setup (see Data Availability and Github). Pipeline execution is controlled via run profiles supporting both HPC environments (recommended) and local executions. These profiles define the pipeline config file, and system-dependent settings, such as job scheduler (e.g. SLURM or PBS) and resource allocation, which can be adapted to the target infrastructure. After configuration, ECHO can be run using a single Snakemake command specifying the relevant execution profile. To facilitate reproducible execution and user validation, ECHO is accompanied by two test ONT datasets: a lightweight smoke-test dataset and a whole-genome dataset (HG002 15×). Both datasets, together with corresponding expected outputs are hosted on Zenodo (see Data Availability).

## 4 Performance evaluation

To benchmark pipeline performance, we applied ECHO to publicly available Genome in a Bottle (GIAB) HG002 ONT data (see Data Availability), basecalled on a PromethION 24 compute tower using Dorado v0.8.3 with the sup_v5,5mCG_5hmCG model. Reads were subsequently subsampled to approximately 30× and 15× genomic coverage to evaluate performance across different sequencing depths. ECHO was run on a HPC system using default configuration settings with the GRCh38 reference build and bundled repeat catalogues (Supplementary Figure 1).

The number of TR loci genotyped by LongTR and the number of detected non-ref-TE insertions passing default TLDR filtering criteria were compared between the 30× and 15× datasets. A moderate reduction in detected loci was observed at lower coverage (Figure 1B), although substantial repeat and methylation calling concordance between datasets was evident (Supplementary Figure 2).

In addition, DNA methylation measurements obtained with ECHO were benchmarked against publicly available HG002 whole-genome bisulfite sequencing (WGBS) data, which is considered the gold standard for genome-wide DNA methylation profiling (Singer, 2019). CpG accuracy was evaluated at three levels: genome-wide, within TEs, and within TRs, defined by the complete GRCh38 TE catalogue and the genome-wide Adotto TR catalogue, respectively. For all datasets, CpG sites with read coverages below 10× or above 200× were excluded to reduce the impact of unreliable measurements due to low coverage and/or misalignment.

For the 30× coverage sample, we observed high concordance with WGBS, as shown via Pearson correlation coefficients of 0.96 (n CpGs = 25,121,849), 0.95 (n CpGs = 12,849,199), and 0.94 (n CpGs = 195,317) for genome-wide, TE, and TR regions, respectively (Figure 1C). As expected, the HG002 15× sample yielded slightly reduced accuracy and fewer CpG sites but still yielded comparable and robust results (Supplementary Table 1). Overall, ECHO achieved DNA methylation accuracy at repetitive regions comparable to genome-wide measurements, indicating it can handle complex regions. Moreover, Integrative Genomics Viewer (IGV) views at representative TE and TR loci support ECHO’s haplotype-resolved integration of repeat genotyping and DNA methylation (Supplementary Figures 3-5).

Finally, the total wall-clock compute time aggregated across the workflow, from basecalled UBAM files to unified haplotype-resolved repeat genotyping and DNA methylation outputs, was 38.5 hours (234 CPU-hours) for the HG002 30× dataset and 26.6 hours (172 CPU-hours) for the 15× dataset. Excluding intermediate preprocessing (UBAM/FASTQ/unphased BAM) and log files, the corresponding ECHO project directories collectively occupied 100 GB (30×) and 60 GB (15×) disk space, respectively.

## 5 Conclusion

We have developed ECHO, a portable, reproducible, and scalable Snakemake pipeline for comprehensive (epi)genomic characterization of the human repeatome using ONT sequencing. By harmonising existing alignment, variant calling, phasing, repeat genotyping and methylation analysis tools within a single workflow, ECHO enables integrated and haplotype-resolved analysis of (epi)genetic variation across multiple classes of repetitive DNA elements, which traditionally have been analysed using separate, specialized approaches so far. Its modular design allows straightforward adjustment, extension, and integration of additional analysis components, ensuring long-term flexibility and compatibility with future tools and updates. Overall, we expect that ECHO will enable and accelerate human repeatome research in the near future at both individual and population level.

## Acknowledgements

The authors thank Nicole Flack and Henry Barton (GeneVia Technologies, Tampere, Finland) for their valuable bioinformatics support and contributions. We also acknowledge Leonard Tiling for his contributions to the development of this pipeline. We further thank Bart de Koning and the Bioinformatics team of the department of Clinical Genetics, Maastricht UMC+ for kindly reviewing the ECHO pipeline documentation and for their support with HPC infrastructure.

## Author contributions

Brando Poggiali: Conceptualization [supporting], Methodology [co-lead], Software [lead], Formal analysis [lead], Visualization [supporting], Writing – original draft [co-lead], Writing – review and editing [supporting]; Leena Putzeys: Conceptualization [supporting], Methodology [co-lead], Software [supporting], Formal analysis [supporting], Supervision [supporting], Visualization [lead], Writing – original draft [lead], Writing – review and editing [co-lead]; Jeppe D. Andersen: Writing – review and editing [supporting]); and Athina Vidaki (Conceptualization [lead], Supervision [lead], Methodology [supporting], Funding acquisition [lead], Writing – review and editing [lead]).

## Supplementary data

Supplementary data is available at *Bioinformatics* online.

Supplementary Figure 1: Content of config.yaml used for ECHO benchmarking.

Supplementary Figure 2: Concordance of repeat calling and methylation profiling between HG002 datasets at 15× and 30× coverage.

Supplementary Figure 3: Haplotype-resolved (epi)genotyping of tandem repeat (TR) with ECHO.

Supplementary Figure 4: Haplotype-resolved (epi)genotyping of reference transposable element (ref-TE) with ECHO.

Supplementary Figure 5: Haplotype-resolved (epi)genotyping of non-reference transposable element (non-ref-TE) with ECHO.

Supplementary Table 1: Concordance between ECHO- and WGBS-derived DNA methylation estimates across genomic contexts and sequencing depths for HG002.

## Data Availability

The pipeline, including documentation and usage guidelines, is available at: https://github.com/leenput/ECHO-pipeline. Any questions, bug reports, and feature requests can be submitted via GitHub.

ONT HG002 dataset:

s3://ont-open-data/giab_2025.01/flowcells/HG002/PAW70337/pod5; WGBS HG002 dataset:

s3://ont-open-data/gm24385_mod_2021.09/bisulphite/cpg/CpG.gz.bismark.cov.gz;

ECHO bundled repeat catalogs:

https://zenodo.org/records/18172438;

smoke test data set and HG002 15× example outputs:

https://doi.org/10.5281/zenodo.19048904; GRCh38 reference genome:

https://ftp.ncbi.nlm.nih.gov/genomes/all/GCA/000/001/405/GCA_000001405.15_GRCh38/seqs_for_alignment_pipelines.ucsc_ids/GCA_000001405.15_GRCh38_no_alt_analysis_set.fna.gz

