## Supplementary data for "ECHO: a nanopore sequencing-based workflow for (epi)genetic profiling of the human repeatome"

#### Content

1. Supplementary Figures ........................................................................ 2
2. Supplementary Tables ........................................................................ 7

### 1. Supplementary Figures


```
samples:
- HG002_30X
- HG002_15X
start_from: ubam
input_dir: ../projects/ECHO_final_benchmark
output_dir: ../projects/ECHO_final_benchmark
reference: ../references/GRCh38/GCA_000001405.15_GRCh38_no_alt_analysis_set.fasta
reference_name: GRCh38
fastq_filtering:
  min_read_quality: 7
  min_read_length: 500
flanking_length_bp: 250
extension_repeat_consensus: 1000
reference_TE: resources/echoDB_v1/TEs/teref.ont.human.fa
type_of_tr: genome-wide
type_of_te: all
tr_catalog: resources/echoDB_v1/TRs/GRCh38/genome-wide/adotto_longTR.bed
te_catalog: resources/echoDB_v1/TEs/GRCh38/GRCh38_TEs_all.bed
tr_methylation:
  filter_to_cpg_str: true
  cpg_str_bed: resources/echoDB_v1/TRs/GRCh38/genome-wide-str
cpg/adotto_longTR_STR_cpgmotif.bed
```


**Supplementary Figure 1: Content of config.yaml used for ECHO benchmarking.**

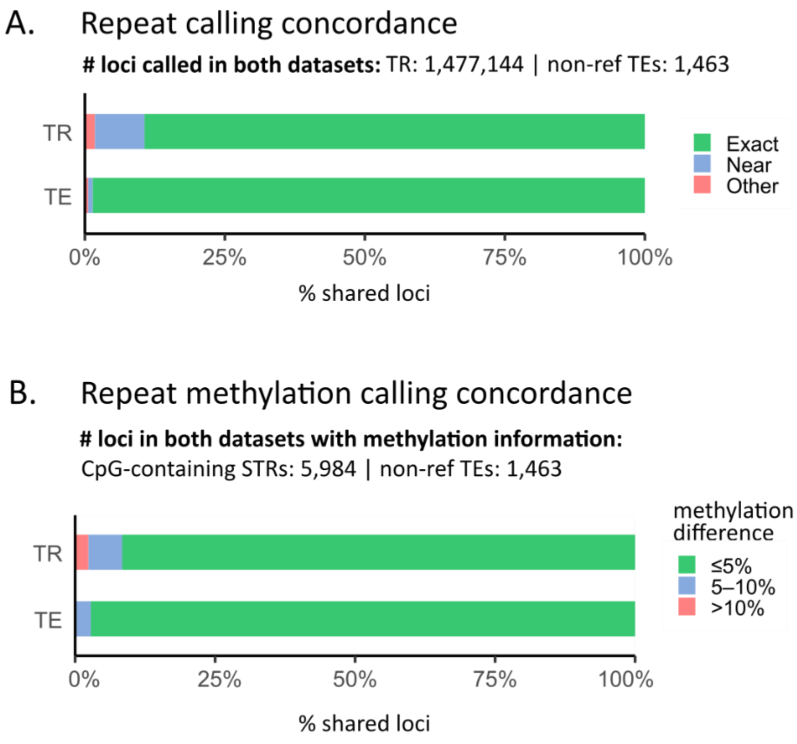


**Supplementary Figure 2: Concordance of repeat calling and methylation profiling between HG002 datasets at 15× and 30× coverage. (A)** Repeat calling concordance was evaluated among loci detected in both datasets by LongTR and TLDR. For tandem repeats (TRs), concordance was assessed across shared TR loci (n = 1,477,144), with loci classified as exact matches (green, identical TR allele lengths), near matches (blue, ≤2 bp difference), or mismatches (red, other discrepancies). For non-ref-TEs, concordance was assessed across shared insertions matched by genomic coordinates and TE family (n = 1,463). Exact matches additionally required concordant insertion lengths (green, ≤5 bp difference), while near matches allowed insertion length differences up to 10 bp (blue). **(B)** DNA methylation concordance was evaluated among shared loci with methylation estimates in both datasets. For TRs, methylation analysis was restricted to loci from the default Adotto TR catalogue whose canonical repeat motif corresponds to a short TR (STR; 1–6 bp) and contains at least one CpG site (n = 8,424). Methylation concordance was assessed among shared CpG-containing STR loci with methylation estimates in both datasets (n = 5,984). For each locus, absolute differences in average per-allele methylation between the 15× and 30× datasets were computed, allowing haplotype label swapping (HP1/HP2). Loci were classified into ≤5%, 5–10%, or >10% difference bins. For non-ref-TEs, methylation concordance was evaluated among shared insertions with methylation measurements for at least one allele in both datasets (n = 1,391), using the same classification thresholds.

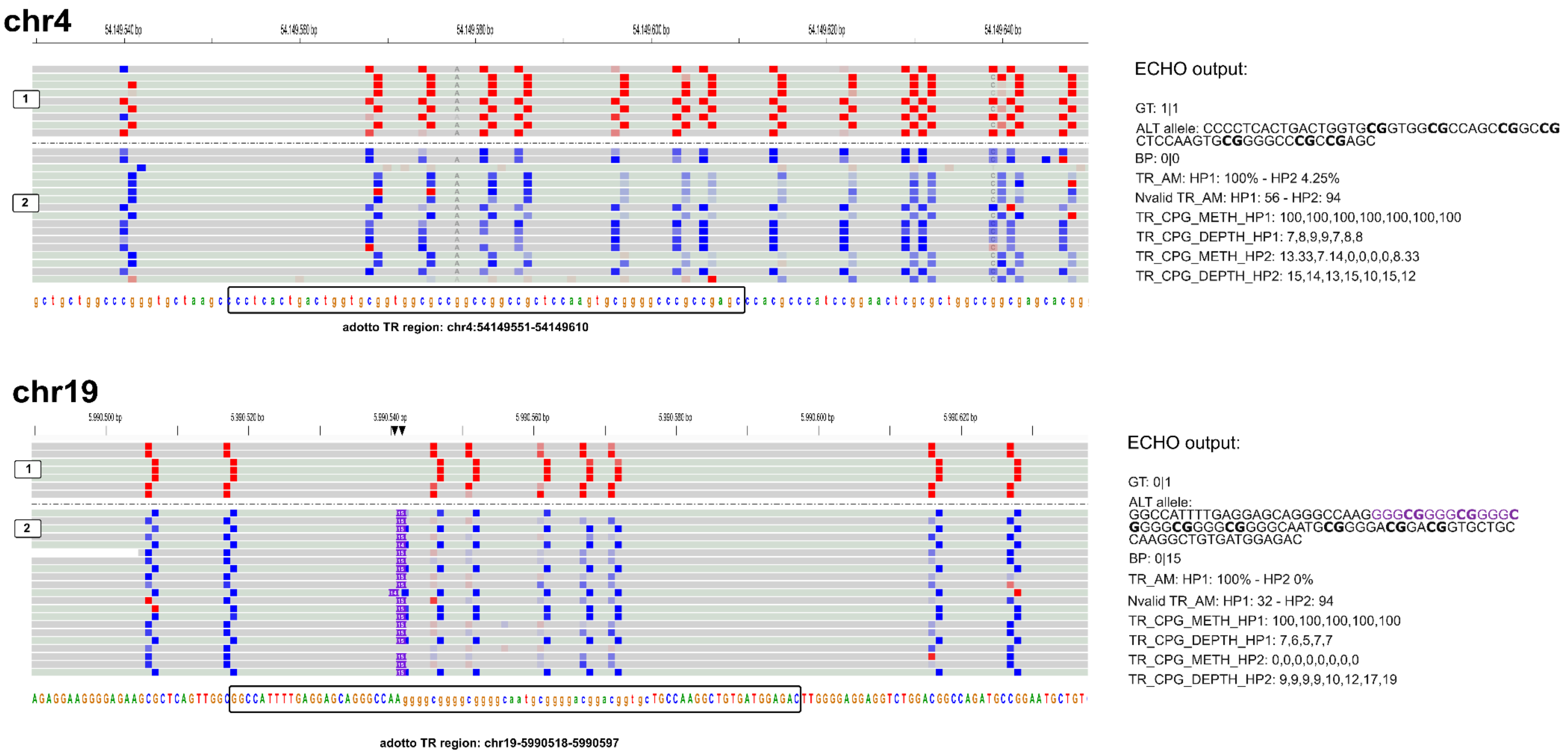


**Supplementary Figure 3: Haplotype-resolved (epi)genotyping of tandem repeats (TR) with ECHO.** IGV snapshots of two example TR loci, as defined by the Adotto catalogue (English et al., 2025) on chr4 (top) and chr19 (below), processed using the ECHO pipeline. Reads were phased into haplotypes (1, 2). Colored blocks indicate CpG methylation states inferred from ONT data (red: methylated; blue: unmethylated). For each TR locus, ECHO reports the TR genotype (GT), inferred alternative (ALT) allele sequence(s), repeat length relative to the reference (BP), and haplotype-specific average methylation levels across the TR region (TR_AM), together with the number of valid methylation observations (Nvalid_TR_AM), and per-CpG methylation (TR_CPG_METH) and depth (TR_CPG_DEPTH) values for each haplotype.

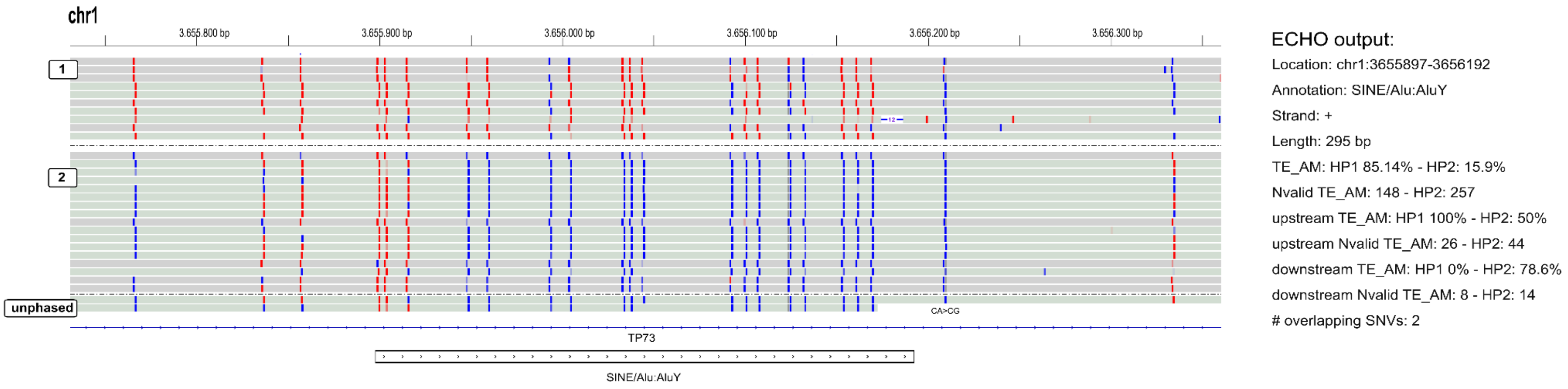


**Supplementary Figure 4: Haplotype-resolved (epi)genotyping of reference transposable elements (ref-TE) with ECHO.** IGV snapshots of an example TE locus (chr1:3,655,897-3,656,192, classification: SINE/AluY), as defined by the UCSC GRCh38 RepeatMasker annotation and processed using the ECHO pipeline. Reads were phased into haplotypes (1, 2). Colored blocks indicate CpG methylation states inferred from ONT data (red: methylated; blue: unmethylated). For each ref-TE, ECHO reports genomic coordinates, TE annotation, repeat length, and haplotype-specific average methylation levels across the reference TE region (TE_AM) and its flanking regions (-250 bp upstream and +250bp downstream), together with the number of valid methylation observations (Nvalid), and the number of overlapping sequence variants, stratified by single nucleotide variants (SNVs) and structural variants (SVs).

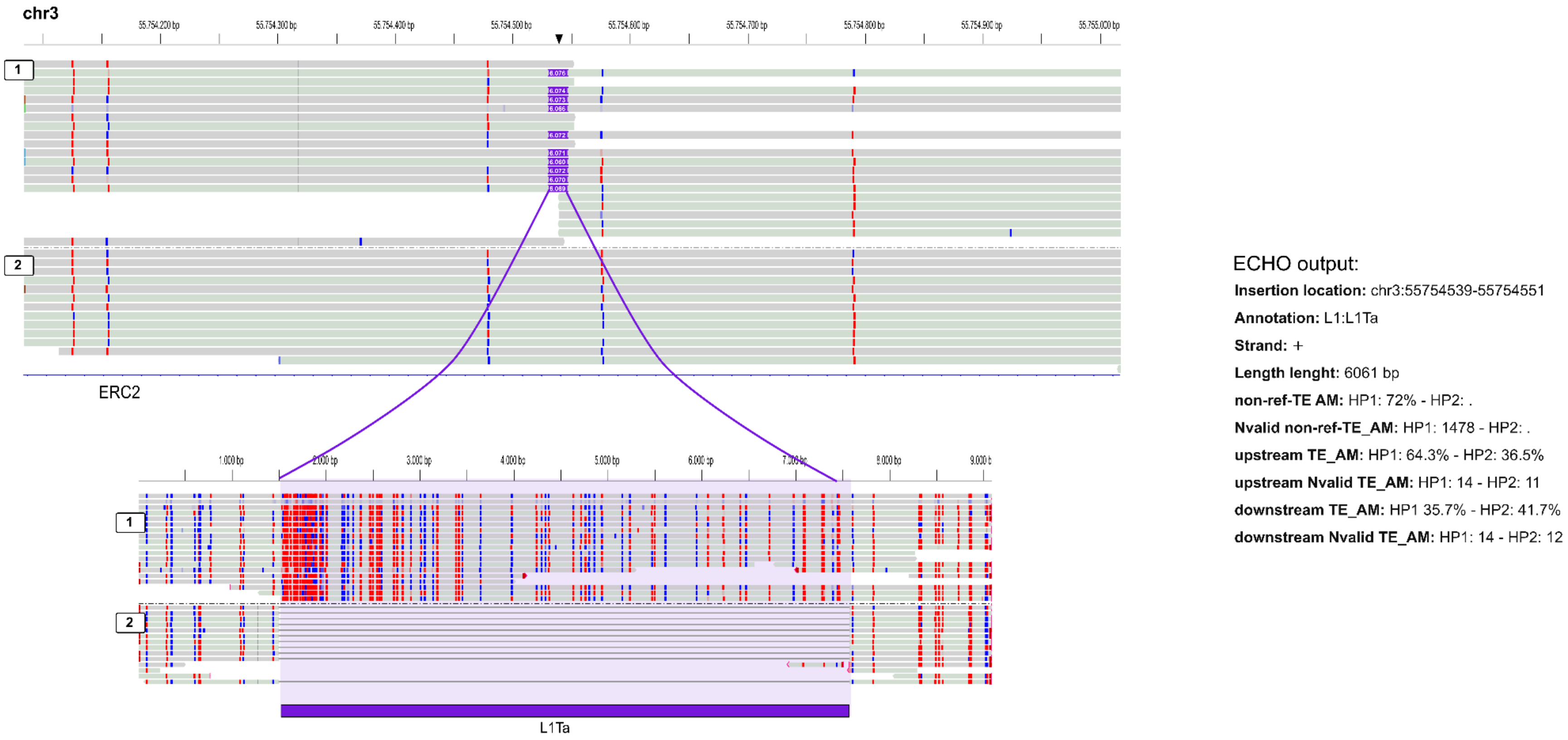


**Supplementary Figure 5: Haplotype-resolved (epi)genotyping of non-reference transposable elements (non-ref-TE) with ECHO.** IGV snapshots of an example non-ref-TE locus (chr3:55,754,539-55,754,551, classification: LINE/L1Ta) detected by TLDR (Ewing et al, 2020) and processed using the ECHO pipeline. Reads were phased into haplotypes (1, 2). Colored blocks indicate insertion events (purple) and CpG methylation states inferred from ONT data (red: methylated; blue: unmethylated). For each non-ref-TE, ECHO reports genomic coordinates, TE annotation, repeat length and sequence, and haplotype-specific average methylation levels across the non-ref-TE insertion (TE_AM) and its flanking regions (-250 bp upstream and +250bp downstream), together with the number of valid methylation observations per haplotype (Nvalid).

#### 2. Supplementary Tables

**Supplementary Table 1: Concordance between ECHO- and WGBS-derived DNA methylation estimates across genomic contexts and sequencing depths for HG002.** Pearson correlation, mean absolute error (MAE), and root mean square error (RMSE) between ECHO- and whole-genome bisulfite sequencing (WGBS)-derived DNA methylation values are shown for CpG sites genome-wide and within transposable elements (TEs) and tandem repeats (TRs). TE and TR annotations were defined using GRCh38 repeat catalogues derived from the UCSC Genome Browser RepeatMasker track and the Adotto TR benchmark (English *et al.*, 2025). Results are shown for ONT HG002 datasets subsampled to 30× and 15× coverage. CpG sites with read coverage below 10× or above 200× in either dataset were excluded.

| **Dataset** | **Context** | **# CpGs** | **Pearson** | **MAE (β)** | **RMSE (β)** |
|---|---|---|---|---|---|
| HG002 30× | Genome | 25,121,849 | 0.9611 | 0.064801 | 0.099676 |
| | TE | 12,849,199 | 0.9515 | 0.064126 | 0.097747 |
| | TR | 2,621,666 | 0.9530 | 0.061877 | 0.100660 |
| HG002 15× | Genome | 20,184,886 | 0.9519 | 0.072843 | 0.106540 |
| | TE | 10,761,518 | 0.9436 | 0.071802 | 0.104314 |
| | TR | 2,074,545 | 0.9410 | 0.067154 | 0.102797 |
